# Marine invertebrate diversity and distribution; the evidence of human activities in Rommel Bay

**DOI:** 10.1101/2022.01.23.477416

**Authors:** Marwa Abd El-Aziz M. Ibrahim, Islam Mohammad Zidan

## Abstract

The current note hypothesised that human/tourism beach activities in Rommel Bay, Marsa Matrouh, Egypt (31°21’39.9”N 27°15’12.3”E) affected the marine invertebrates in this area. The note resulted some new findings of arthropod, molluscan, and seagrasses species diversity and distribution. The hypothesised study concluded that human/tourism beach activities did not eliminate the invertebrate communities, but these animals have shifted and moved themselves to escape away from noise and pollution. Further investigations should be taken into consideration about aquatic insects, mites, crustaceans, seagrasses, and molluscans diversity and distribution, to figure out the actual structures and interactions between these animals within the Rommel Bay ecosystem, besides what are the stresses they have, and what are the consequences.

## Introduction

The North-western coast of Egypt, from Alexandria to El-Salum, is about 510 km (Tadros *et al.*, 2015). According to Egypt’s Prime Minister report; the tourism sector in Marsa Matrouh reached 43 hotels and resorts (about 8500 beds) in 2019 hosted more than seven million visitors in 2017 during summer months (May to Sep) (IDSC, 2021). The coastal line of Matrouh is protected by rock walls from the open sea with some openings to allow water to renew and flow. Four bays/lagoons formed due to these rocky walls; Al-Gharam Bay (Cleopatra), Matrouh Bay, Rommel Bay, and Alam Al-Rom Bay (Al-Fairouz) (Gharib *et al.*, 2011; Abdel Ghani, 2015; Elhmamy *et al.*, 2019) (Figure 1 top).

**Figure 1.**
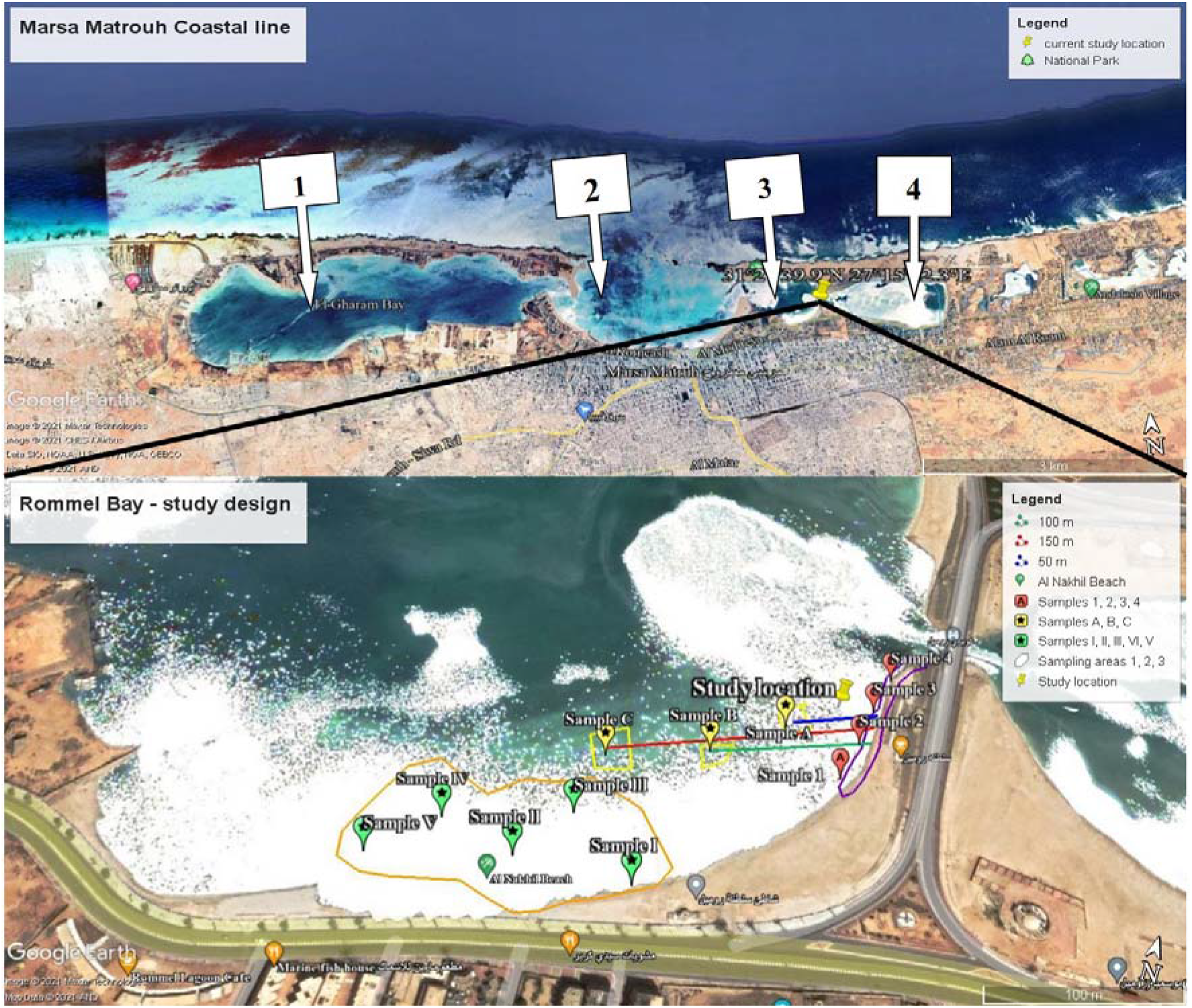
**(Top)** Bays/lagoons of Matrouh; 1) Al-Gharam Bay (Cleopatra), 2) Matrouh Bay, 3) Rommel Bay, and 4) Alam Al-Rom Bay (Al-Fairouz). **(Bottom)** Study location, where the purple line representing the rocky shore (area 1) which consisted of four samples (red drop pins); the yellow drop pins representing area 2 and the length of samples; 50 m (blue line), 100 m (green line), and 150 m (red line). The orange area with green drop pins has five different samples (area 3).

Integration of the abiotic factors as physical (e.g., transparency, dissolved oxygen, salinity, sea level, and pH) and the chemical properties, (e.g., NO3, NO2, NH4, PO4, and SiO2), and the biotic factors limits the diversity and distribution of marine organisms (Azov, 1991; Frihy, 1992; Gharib *et al.*, 2011; Lhullier *et al.*, 2020). Also, the climatic changes have affected the heat storage values of water column (J/m^2^) of the Mediterranean basin in the upper 100 and 300 m layers, when reached their highest levels from Aug to Oct and the lowest levels from Feb to Apr annually (Kamel *et al.*, 2020). As well, the water temperature from nine beaches of Matrouh has been detected, 17.45-18.40 °C in winter and 27.25-32.00°C in summer (Gharib *et al.*, 2011). Due to these factors; the phytoplankton diversity indices varied between 83.75% (Bacillariophyta group which represented by 61 genera and 120 species) to 0.04% (Silicoflagellates group which represented by only one genus and species) during four seasons in nine beaches of Matrouh (Gharib *et al.*, 2011). Thus, this short note proposed evidence of the tourism/human activities that affected the diversity and distribution of the marine invertebrates in Rommel Bay, Marsa Matrouh to study these hypothesised indices in the future.

## Materials and Method

To test the theory, samples have been collected from Al Nakhil Beach, Rommel Bay, Marsa Matrouh (31°21’39.9”N 27°15’12.3”E). We divided the location into three sampling areas, as shown in Figure (1 bottom); i) area 1 is four samples of the rocky coastal shore; ii) area 2 containing three samples of 50, 100, and 150 m away from the shore (length measured by GPS using Google maps and Google Earth) and 1.5 m in depth underwater; iii) area 3 is five samples of different sites of the beach and beneath the sea (1 – 1.5 m in depth).

Biodiversity Shannon-Wiener index (H’), and Simpson’s index (dominance D and species richness 1/D) were calculated for all samples using the BioDiversity Pro. ver. 2.0 software (McAleece *et al.*, 1997) and PAST ver. 4.03 software (Hammer *et al.*, 2001). Differences in the mean number of species within and/or among the areas were evaluated using one-way analysis of variance (ANOVA) and were tested with Tukey’s test at 95% confidence level using SPSS computer program ver. 20.0.

## Results and Discussion

There was a statistically significant difference in the diversity and distribution of the samples (*F*= 2.46, *df*= (4, 60), *P*= 0.03) at confidence level of 95%. There was a significant difference between sample medians that tested using Kruskal-Wallis; *H*_*0*_= 14.72, *H*_*1*_= 18.79, *P*= 0.000 at the same confidence level. Various kinds of groups were connected in a complexed structure with a lot of similarities and dissimilarities (Figure 2. a & b), which theoretically proposed a complex food web. Seagrass, *Posidonia oceanica* (L.), has dominantly existed in all samples of the location. However, it was distantly different from area 1 to area 3, when was intensively gathered from the rocky shore than from places that were full of human activities (D= 0.259, 1/D= 3.849, H’= 1.440). The role of *P. oceanica* has been revealed as a pollution bio-indicator in the Mediterranean basin (Telesca *et al.*, 2015).

**Figure 2.**
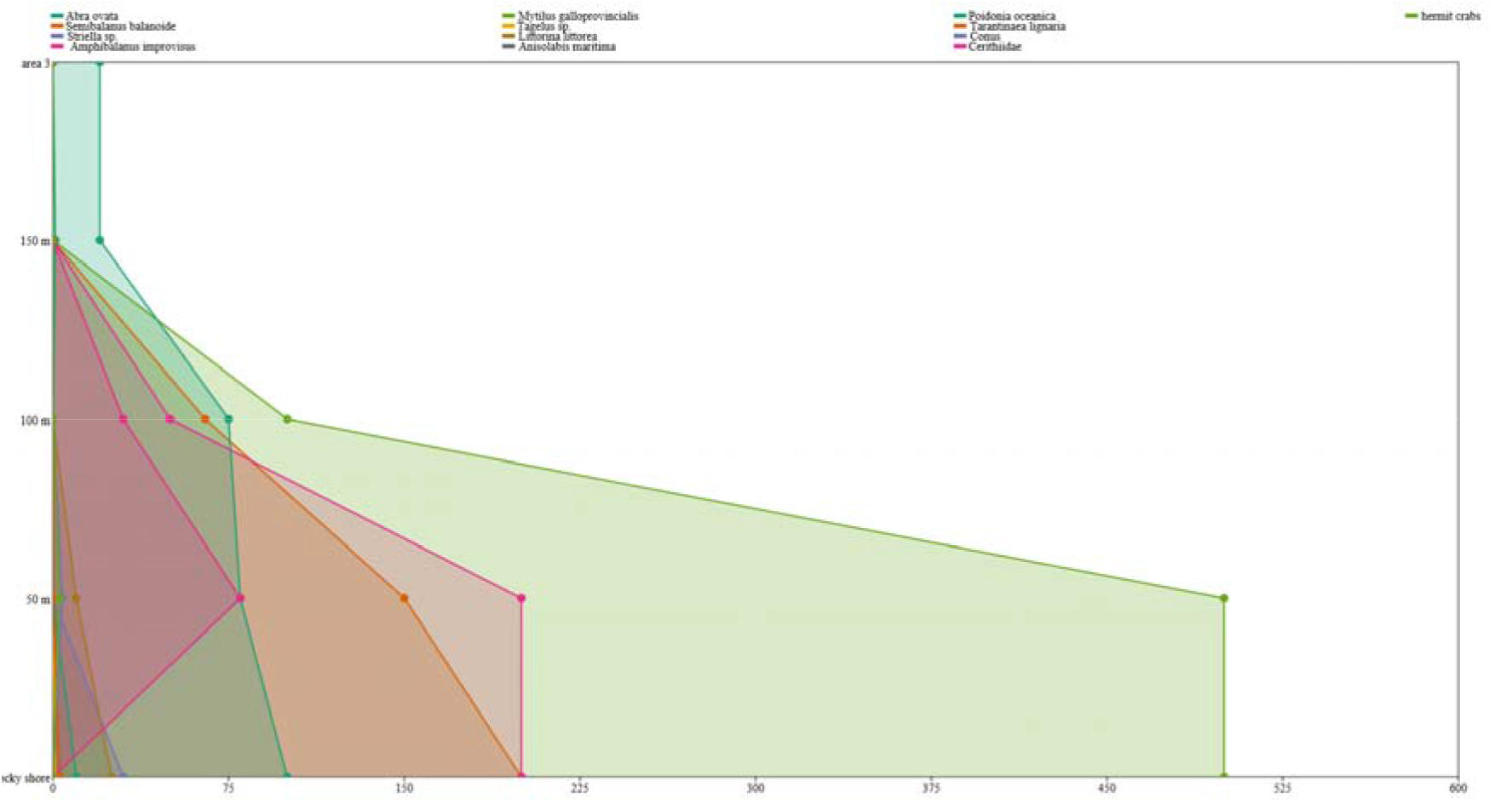
**a**. Species diversity indices and distribution in each area

**Figure 2.**
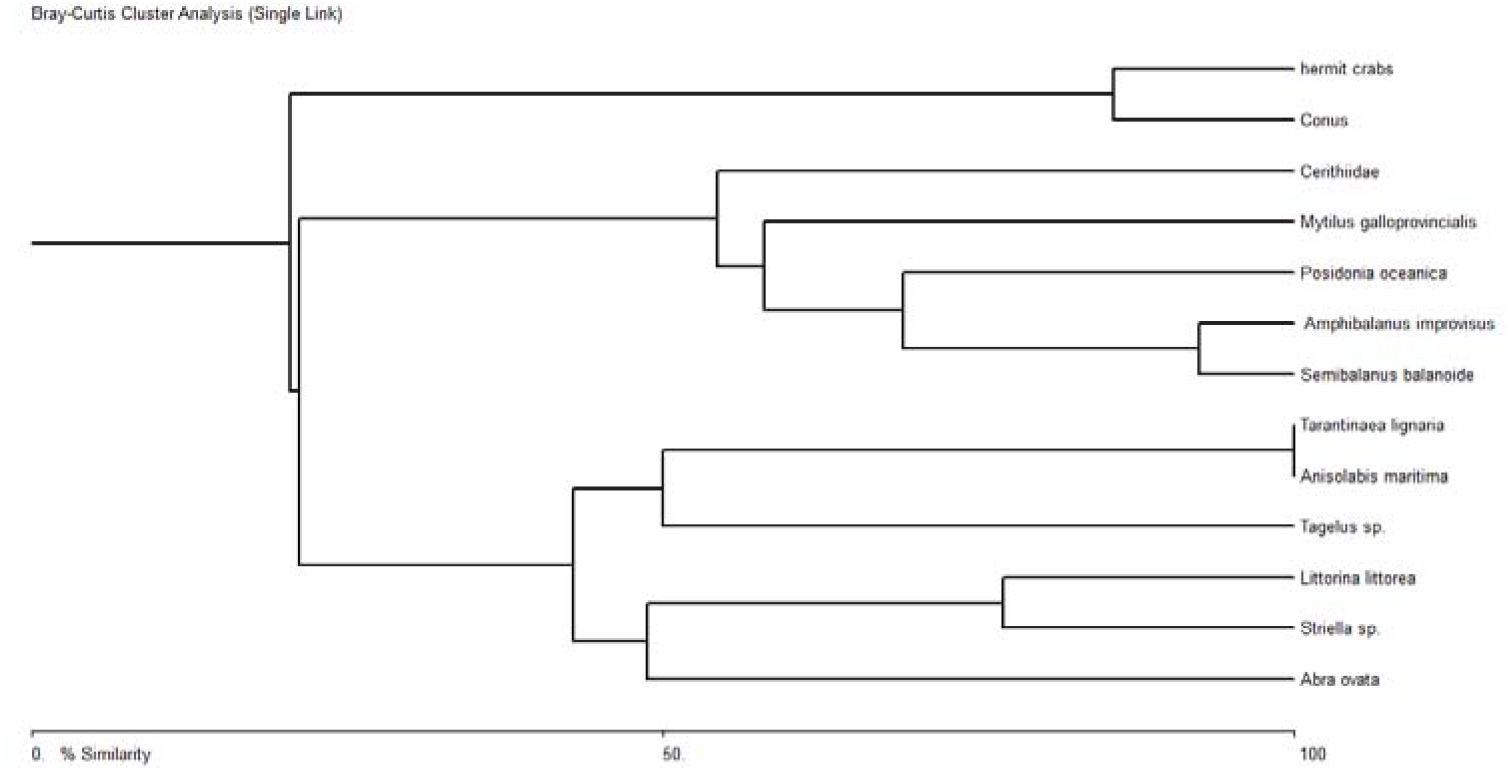
**b**. Species distribution similarity and dissimilarity within Rommel Bay experimental location. The rocky shore (area 1) had the highest diversity indices recorded comparing with other sampling areas within Rommel location (D= 0.297, 1/D= 3.372, H’= 1.478). Although, this side was containing the maximum number of individuals and taxa (S) sampled, but there were no significant differences detected among these taxa (total individuals= 1072, S= 10, *T*= 2.05, *df*= 12. *P*= 0.063) (Figure 3) (Supplementary 1).

The proposed food web occurred in areas 1 & 2 contains a lot of molluscans, crustaceans, and earwigs, when *P. oceanica* was the energy producer and level one food source. Fish deceased bodies, *P. oceanica* remains, and rotten food materials decomposers represented by fungi and some crustaceans which been found beneath rocks, as well have been considered a food source for the different invertebrate species in this area (Figures 3–4).

**Figure 3.**
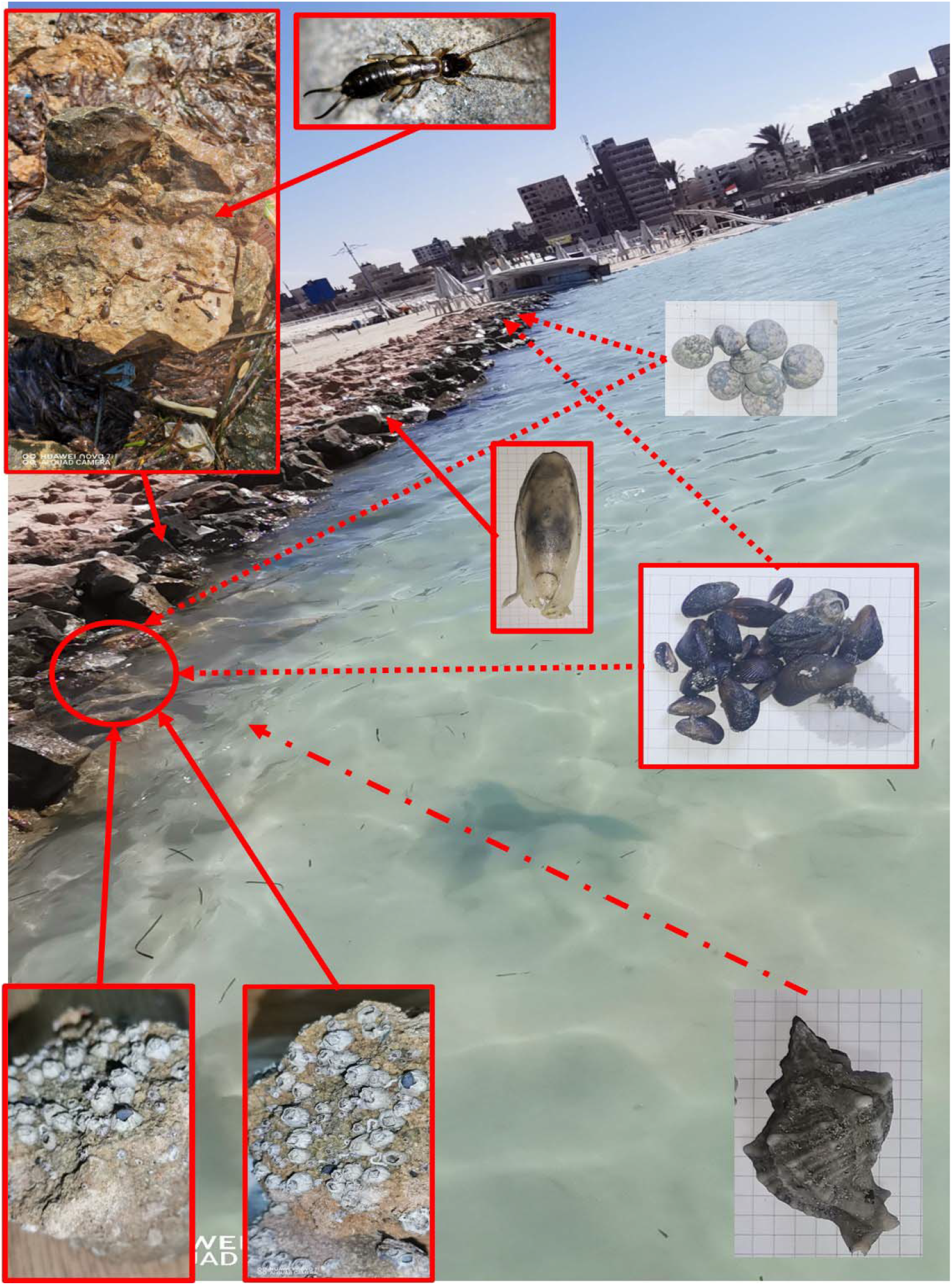
Area 1 recorded huge population of crustacean species that inhabiting rocky shore and beneath water, clusters of seagrasses were over rocks and beneath water, and a lot of human pollution over the rocks along the shore.

**Figure 4.**
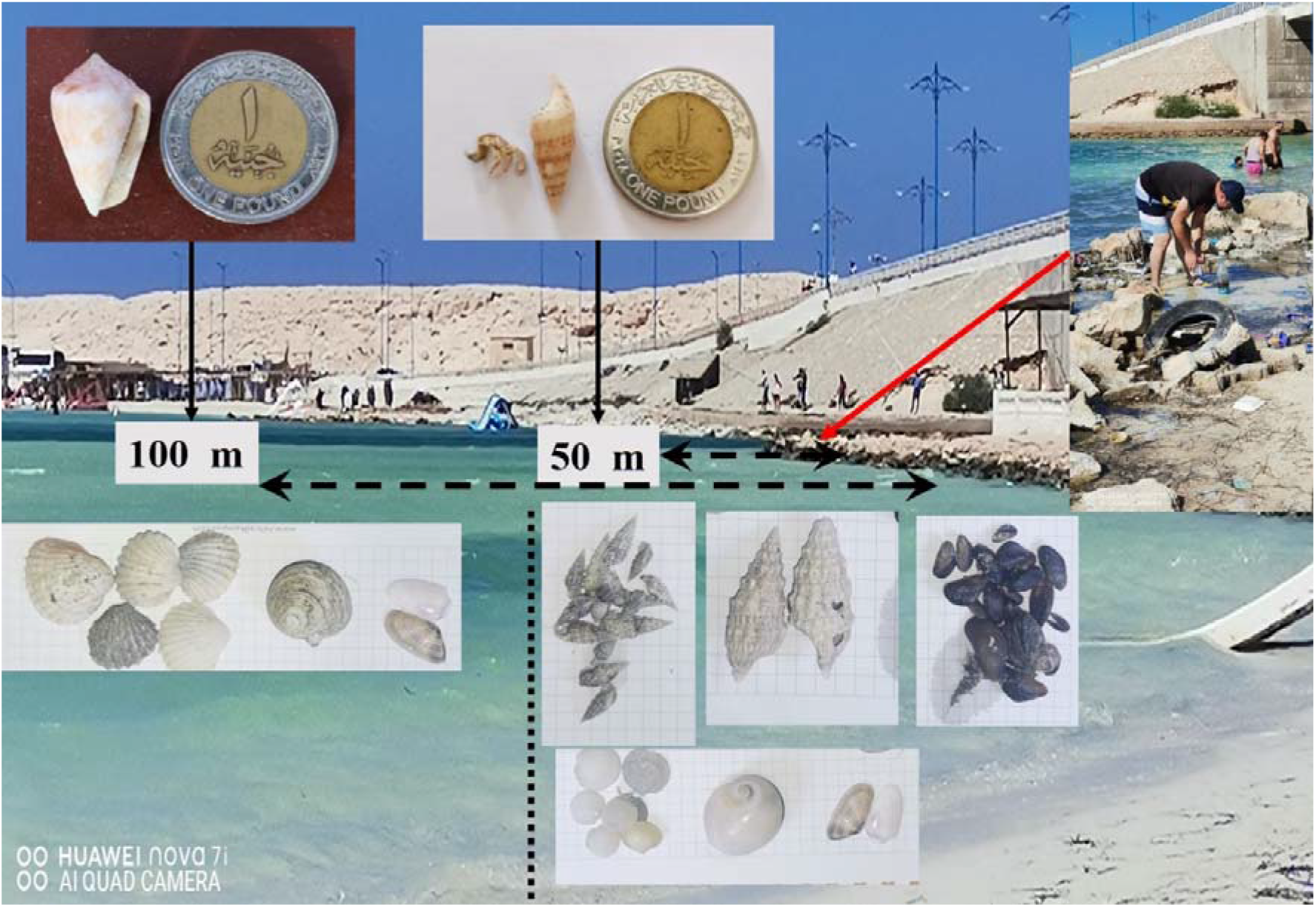
Area 2 has a less species richness and a large mollusc species diversity.

The common barnacle, *Semibalanus balanoides* (Linnaeus, 1767) (D= 0.385, 1/D= 2.591, H’= 1.012), and the bay barnacle, *Amphibalanus improvisus* (Darwin, 1854) (D= 0.406, 1/D= 2.462, H’= 0.967) (Balanidae: Balanomorpha: Crustacea: Arthropoda) were distributed along the rocky shore and the rocks beneath the water in areas 1 & 2 and their distribution has a close similarity (Figure 2.b), nevertheless they were not found on the area 3 that has a lot of human activities. The scavenger *Striella* sp. Glynn, 1968 (Sphaeromatidae: Isopoda: Crustacea: Arthropoda) has no diversity indices (D= 1, 1/D= 1, H’= 0) (Supplementary 2). The null indices were due to this species has only one location had collected from, and it has been known to be found on rocky shores among barnacles (Harrison and Ellis, 1991).

The common periwinkle *Littorina littorea* (Linnaeus, 1758) (Littorinidae: Littorinimorpha: Gastropoda: Mollusca) was found in big density on area 1 (D= 0.579, 1/D= 1.725, H’= 0.612), that might be fed on algae and fungi of rocks or the *P. oceanica* decomposed remains (Figure 3). However, it’s closest similar was *Striella* sp, (Figure 2. b).

Mussels are serving to be good bioindicators of environmental pollution (Sanpanich and Wells, 2019), so that we found massive clusters of the Mediterranean mussel *Mytilus galloprovincialis* Lamarck, 1819 (Mytilidae: Mytilida: Bivalvia: Mollusca) were stuck on shore rocks and/or beneath the water (D= 0.421, 1/D= 2.375, H’= 0.936). An only specimen of the mussel *Tagelus* sp Gray, 1847 (Solecurtidae: Cardiida: Bivalvia: Mollusca) was found on area 1 recorded null indices, while no mussel species were sampled from areas 2 & 3. Few samples of the seaside earwig *Anisolabis maritima* (Bonelli, 1832) (Anisolabididae: Dermaptera: Insecta: Arthropoda) were found beneath rocks, feeding on the *Striella* sp. individuals. We theoretically hypothesised that *A. maritima* representing the predatory role in this complex of organisms, however, the different feeding behaviour of *A. maritima* (Guppy, 1949; Griffiths, 2018), and due to its high dominancy and richness for the area 1 (Figure 3).

Three Mediterranean tulip shell snail *Tarantinaea lignaria* (Linnaeus, 1758) (Fasciolariidae: Gastropoda: Mollusca) have been found, one was alive on the rocky shore, and the other two individuals were broken on the rocks, it was suggested that they have been eaten by a Corvus *Corvus corax* Linnaeus, 1758 (Corvidae: Passeriformes: Aves: Chordata) which was been noticed flying around the place. Another unidentified bird species has been watched, the Bee-eater *Merops* sp Linnaeus, 1758 (Meropidae: Coraciiformes: Aves: Chordata), which is projected to be found there as a resting point on his long journey within the Mediterranean basin. These birds were suggested to be a part of the proposed food web complex, due to their feeding on rock molluscans and arthropods.

Area 2, which has three samples differed in distribution; the 50 m away samples have much diversity and richness (*S*= 8, total individuals= 1027, *D*= 0.308, *1/D*= 3.248, *H’*= 1.435) than those of 100 m (*S*= 5, total individuals= 320, *D*= 0.225, *1/D*= 4.453, *H’*= 1.546) and 150 m (*S*= 2, total individuals= 21, *D*= 0.905, *1/D*= 1.105, *H’*= 0.215), and the conducted results were significantly different of 50 m (*F*= 2.00, *df*= 12, *P*= 0.069), 100 m (*F*= 2.46, *df*= 12, *P*=0.029), and 150 m (*F*= 1.05, *df*= 12, *P*= 0.313) (Supplementary 1).

Measuring samples of 150 m distance (which had been closer to human activities on the beach), there were empty shells of the bivalve mollusc *Abra ovata* (Philippi, 1836) (Semelidae: Cardiida: Bivalvia: Mollusca) barely to count (*D*= 0.818, *1/D*= 1.222, *H’*= 0.350). Unidentified cone snail specimens belonging to the family Conidea, Fleming, 1822 (Gastropoda: Mollusca) (probably genus *Conus* Linnaeus, 1758) were found on a few numbers underwater, 50 m away of the rocky shore, resulting high abundance, species richness, and no diversity indices. While ceriths shells of the family Cerithiidae, Fleming, 1822 (Gastropoda: Mollusca) been found on large numbers 50 m, few numbers on 100 m, and hardly were been found on 150 m in area 2 (*D*= 0.600, *1/D*= 1.668, *H’*= 0.591). Three specimens of the cerithiids have tiny hermit crabs (Crustacea: Arthropoda) inside their spiral shells, 50 m distance (Figure 4) (Supplementary 2).

Samples of area 3 have a low *P. oceanica* existence (Figure 5).

**Figure 5.**
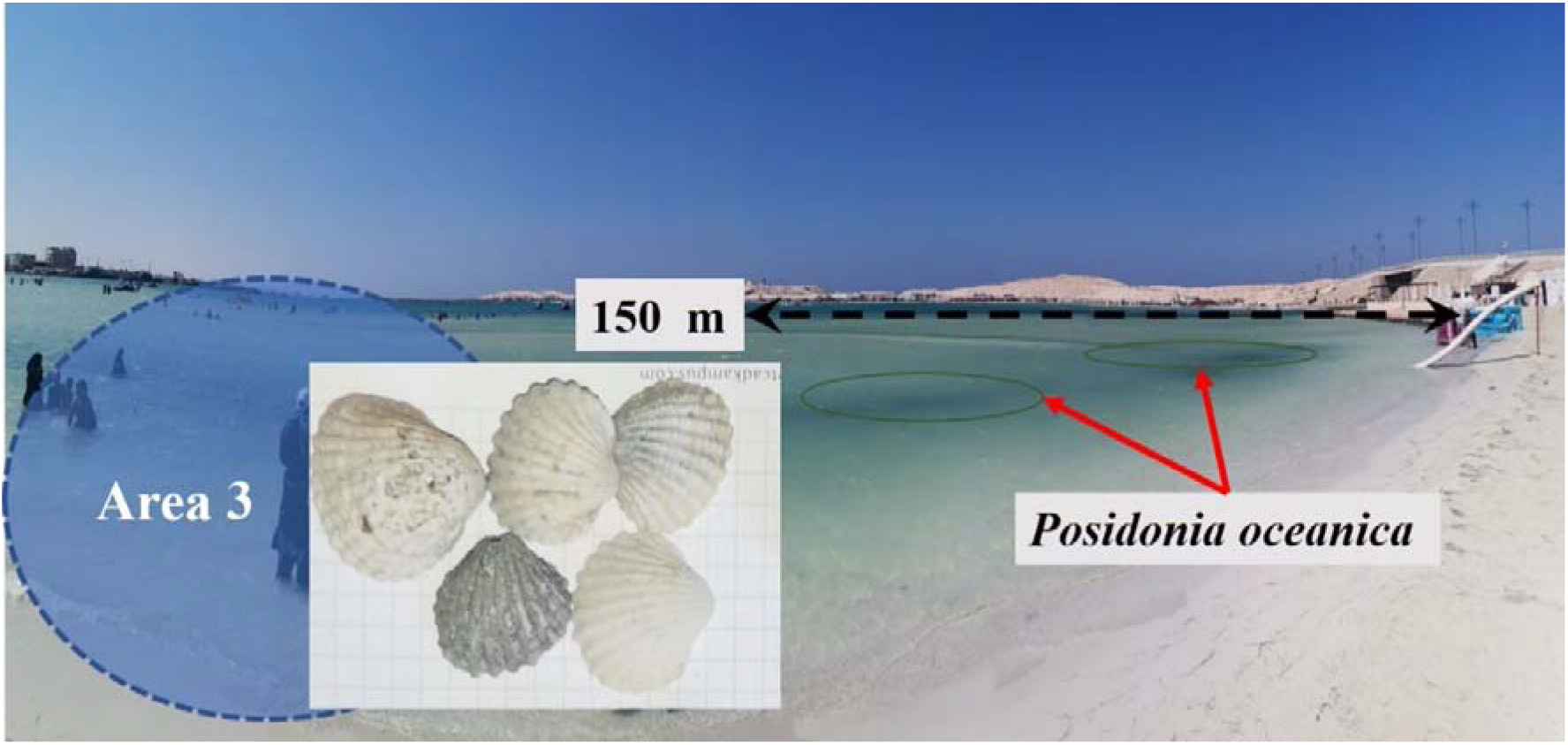
Area 3 has barely few species to be counted. It was predicted to have null diversity indices due to human activities.

It was assumed that plastic bottles, food materials, and human movements significantly affected the invertebrate distribution and diversity in this area. Marine animals that found in different structures during winter when no tourism activities occurred, as the form of the coast has changed because of climatic effects and urban expansions through years (Iskander *et al.*, 2008; Elhmamy *et al.*, 2019; Iskander, 2021). Also, rapid, nonenvironmentally, and inadequately planned coastal projects and activities affected the biodiversity of the marine ecosystem, which requires a necessity of the Environmental Impact Assessment (EIA) (Frihy, 2001; Tabet and Fanning, 2012; Elhmamy *et al.*, 2019; Abdel Kader and Haron, 2020).

## Conclusion

The note suggested that human/tourism beach activities did not eliminate the invertebrate communities, but these animals have shifted and moved themselves to escape away from noise and pollution. Further investigations should be taken into consideration about aquatic insects, mites, crustaceans, seagrasses, and molluscans diversity and distribution, to figure out the actual structures and interactions between these animals within the Rommel Bay ecosystem, besides what are the stresses they have, and what are the consequences.

## Acknowledgments

We would like to express our deep gratitude to Moaaz, Omar, and Farida who helped in collecting samples. Thanks are due to Paula and Dina for their help in the gastropods collection.

## Discloser

The authors declare that there is no conflict of interest regarding the publication of this short communication.

