## Supplementary 1 & 2 for "Marine invertebrate diversity and distribution; the evidence of human activities in Rommel Bay"

| **Supplementary 1. Diversity indices of the Rommel Bay sampling areas** | | | | | |
| --- | --- | --- | --- | --- | --- |
|  | **Area 1 (Rocky shore)** | **Area 2** | | | **Area 3** |
|  |  | **50 m** | **100 m** | **150 m** |  |
| **Total samples** | 13 | 13 | 13 | 13 | 13 |
| **Taxa (S)** | 10.00 | 8.00 | 5.00 | 2.00 | 1.00 |
| **Individuals** | 1072.00 | 1027.00 | 320.00 | 21.00 | 20.00 |
| **Mean** | 82.46 | 79.00 | 24.62 | 1.62 | 1.54 |
| **±SE** | 40.29 | 39.57 | 9.93 | 1.53 | 1.54 |
| **Dominance (D)** | 0.297 | 0.308 | 0.225 | 0.905 | 1.000 |
| **Simpson (1/D)** | 3.372 | 3.248 | 4.452 | 1.105 | 1.000 |
| **Evenness (e^H/S)** | 0.439 | 0.525 | 0.938 | 0.620 | 1.000 |
| **Shannon (H')** | 1.478 | 1.435 | 1.546 | 0.215 | 0.000 |
| ***T*** | 2.05 | 2.00 | 2.48 | 1.05 | 1.00 |
| ***df*** | 12 | 12 | 12 | 12 | 12 |
| ***P*** | 0.063 | 0.069 | 0.029 | 0.313 | 0.337 |

| **Supplementary 2. Diversity indices of the marine invertebrates and seagrass samples** | | | | | |
| --- | --- | --- | --- | --- | --- |
|  | **Individuals** | **Dominance (D)** | **Simpson (1/D)** | **Shannon (H')** | **Evenness (e^H/S)** |
| Abra ovata | 11.00 | 0.818 | 1.222 | 0.350 | 0.710 |
| ***Semibalanus balanoide*** | 415.00 | 0.386 | 2.591 | 1.012 | 0.917 |
| ***Striella sp.*** | 30.00 | 1.000 | 1.000 | 0.000 | 1.000 |
| ***Amphibalanus improvisus*** | 450.00 | 0.406 | 2.462 | 0.967 | 0.877 |
| ***Mytilus galloprovincialis*** | 1100.00 | 0.421 | 2.375 | 0.936 | 0.850 |
| ***Tagelus sp.*** | 1.00 | NAN | NAN | 0.000 | 1.000 |
| ***Littorina littorea*** | 35.00 | 0.580 | 1.725 | 0.613 | 0.923 |
| ***Anisolabis maritima*** | 3.00 | 1.000 | 1.000 | 0.000 | 1.000 |
| ***Posidonia oceanica*** | 295.00 | 0.260 | 3.849 | 1.440 | 0.845 |
| ***Tarantinaea lignaria*** | 3.00 | 1.000 | 1.000 | 0.000 | 1.000 |
| ***Conus*** | 4.00 | 1.000 | 1.000 | 0.000 | 1.000 |
| **Cerithiidae** | 110.00 | 0.600 | 1.668 | 0.591 | 0.902 |
| **hermit crabs** | 3.00 | 1.000 | 1.000 | 0.000 | 1.000 |
| ***df*** | (12, 52) |  |  |  |  |
| ***F*** | 2.989 |  |  |  |  |
| ***P*** | 0.003 |  |  |  |  |

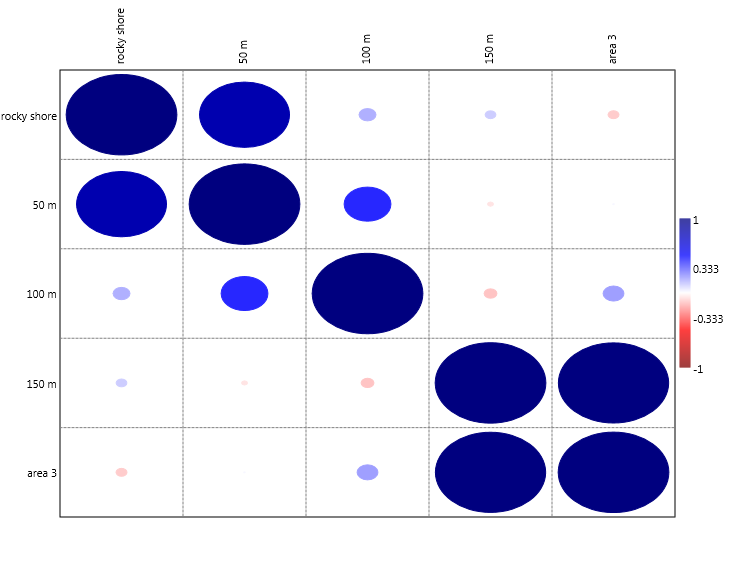

Supplementary 3. Correlation (Partial Linear)

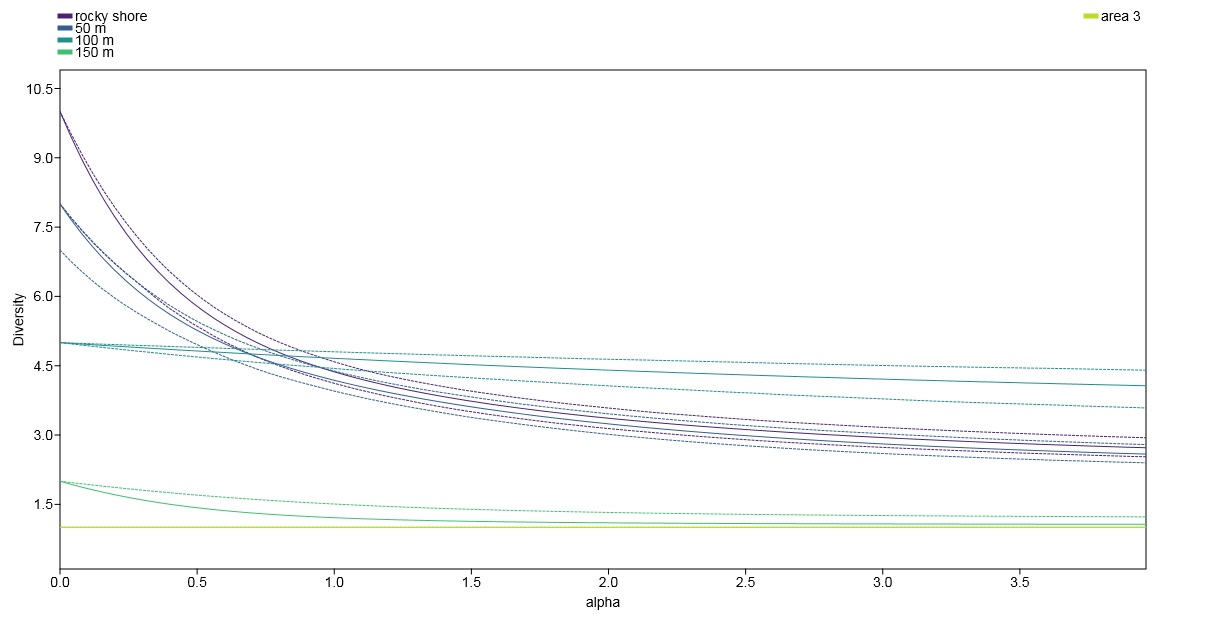

Supplementary 4. Diversity profiles of the Rommel location

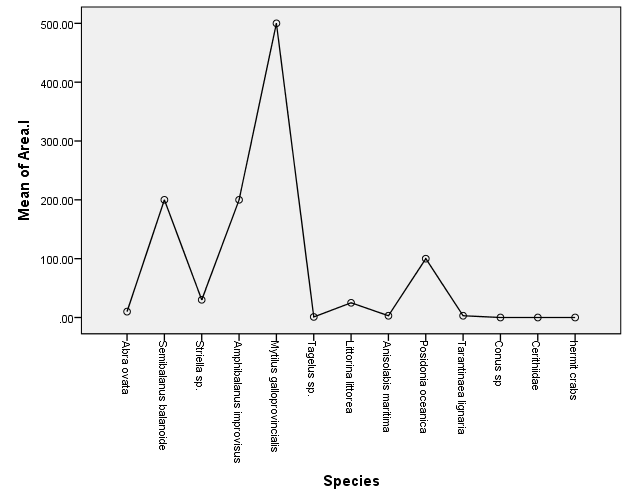
Supplementary 5. Charts of the mean number of speacies in each sampling area

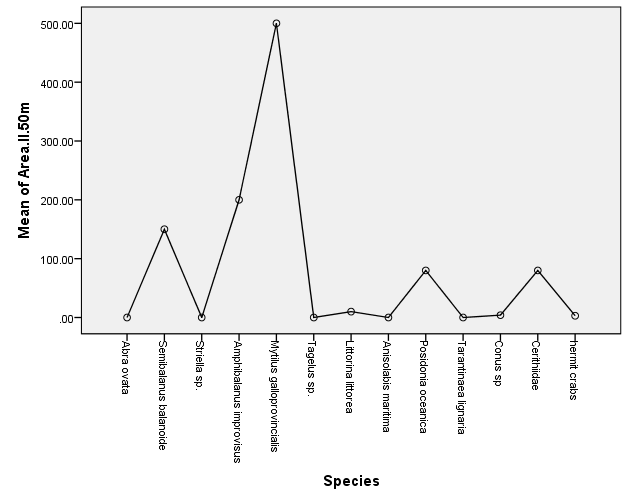

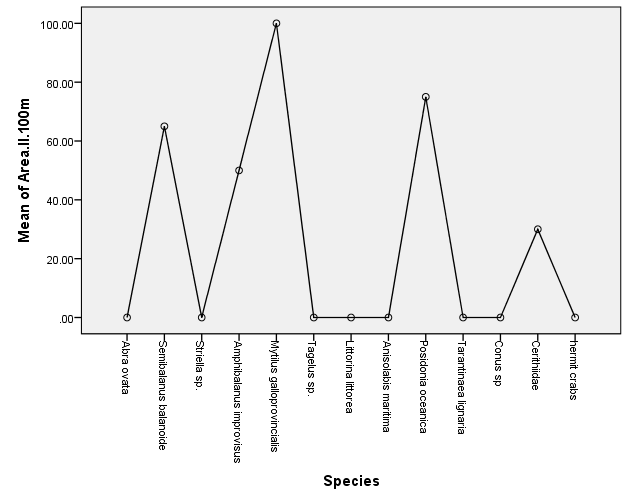

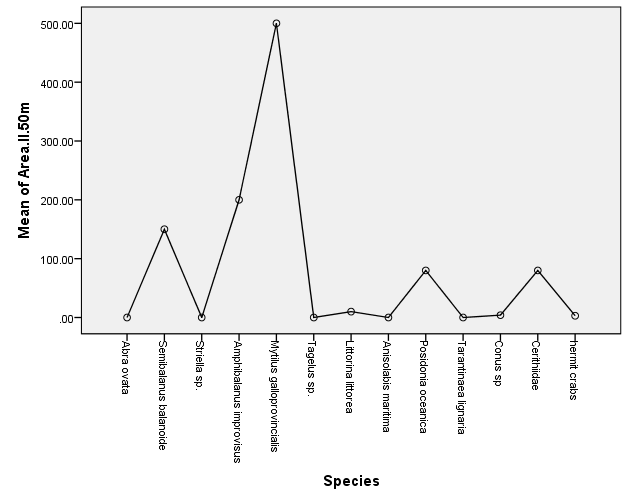

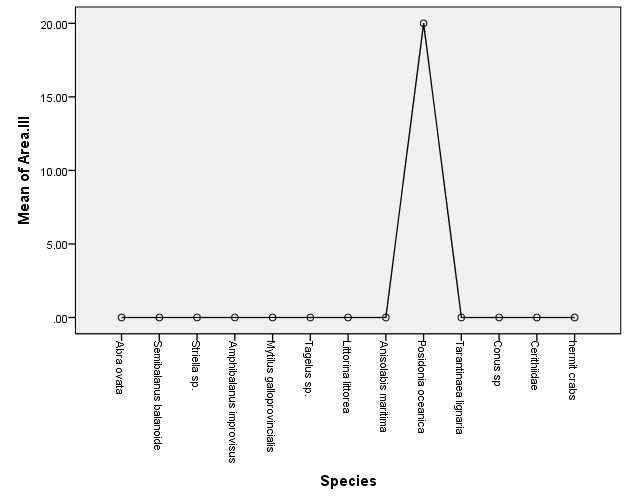

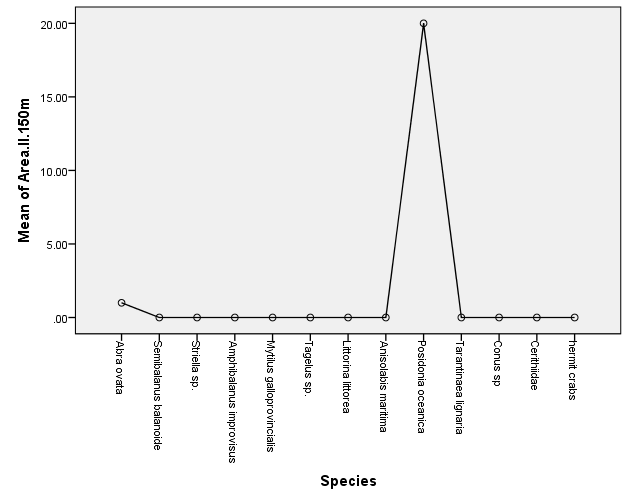

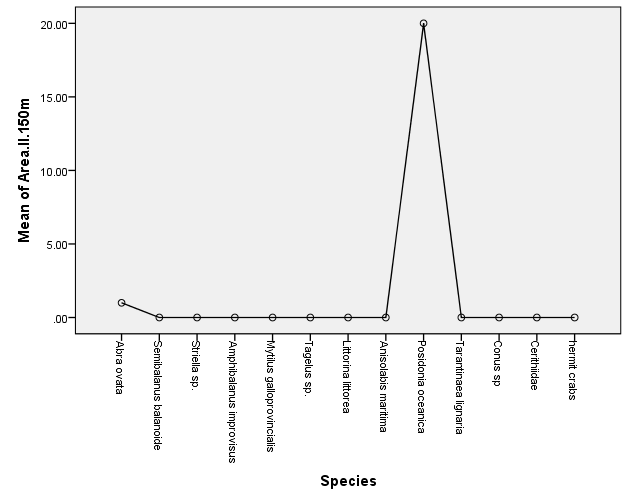

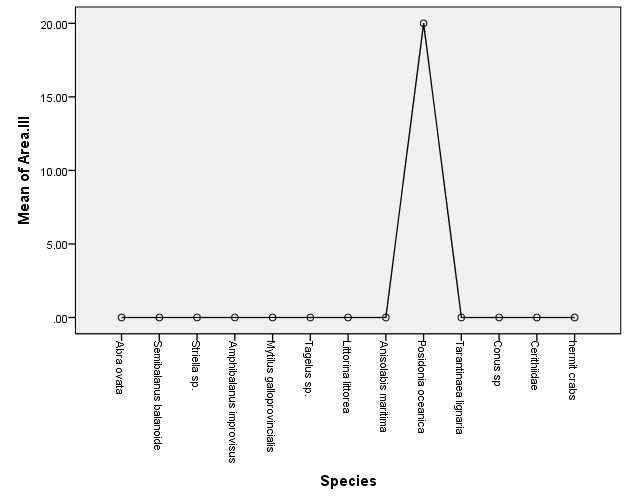
